# The Genome-wide Signature of Short-term Temporal Selection

**DOI:** 10.1101/2023.04.28.538790

**Authors:** Michael Lynch, Wen Wei, Zhiqiang Ye, Michael Pfrender

## Abstract

Despite evolutionary biology’s obsession with natural selection, few studies have evaluated multi-generational series of patterns of selection on a genome-wide scale in natural populations. Here, we report on a nine-year population-genomic survey of the microcrustacean *Daphnia pulex.* The genome-sequences of > 800 isolates provide insights into patterns of selection that cannot be obtained from long-term molecular-evolution studies, including the pervasiveness of near quasi-neutrality across the genome (mean net selection coefficients near zero, but with significant temporal variance about the mean, and little evidence of positive covariance of selection across time intervals), the preponderance of weak negative selection operating on minor alleles, and a genome-wide distribution of numerous small linkage islands of observable selection influencing levels of nucleotide diversity. These results suggest that fluctuating selection is a major determinant of standing levels of variation in natural populations, challenge the conventional paradigm for interpreting patterns of nucleotide diversity and divergence, and motivate the need for the development of new theoretical expressions for the interpretation of population-genomic data.

**Significance:** Except for mono/oligogenic traits known in advance to be under strong selection, there is little information on genome-wide patterns of temporal dynamics of allele-frequency changes in well-defined and unmanipulated natural populations. A multi-year survey of a population of the microcrustacean *Daphnia pulex* provides insight into these matters. Genome-wide analysis of > 800 genetic isolates demonstrates that temporal variation in selection intensity is a major determinant of levels of nucleotide polymorphism and divergence. Most nucleotide sites experience fluctuating selection with mean selection coefficients near zero, with little covariance in the strength of selection across time intervals, and with selection distributed across large numbers of genomic islands of linked sites. These results raise challenges for the conventional interpretation of measures of nucleotide diversity and divergence as indicators of effective population sizes and intensities of positive/negative selection.

---

No biologist needs convincing that natural selection is a powerful force influencing essentially all organisms. Less clear, however, is the magnitude and temporal dynamics of selection operating across genomic regions and across time in natural populations. Hundreds of studies have focused on the measurement of fitness functions associated with quantitative traits, and many more have pursued genome-wide scans of divergent lineages in attempts to pinpoint nucleotide sites under purifying vs. positive selection at individual gene loci (Hughes 2000; Graur 2016; Walsh and Lynch 2018). Whereas the first class of studies yields direct short-term measures at the level of complex traits, the targets of study are generally chosen because they are thought *a priori* to be under a particular form of selection, and the molecular underpinnings are generally left unstudied (Endler 1986; Kingsolver and Diamond 2011). In contrast, whereas the second class of studies is focused at the molecular level, the time scale of divergence is commonly on the order of many thousands to millions of generations, and numerous unobserved aspects of demography and population structure can obscure the interpretation of observed patterns (Johri et al. 2022). Recent studies of evolving laboratory populations have greatly enhanced our understanding of the origin of adaptations at the molecular level, but such studies generally focus on a specific induced selective challenge and/or initiate with a synthetic population structure, including the reliance on the emergence of *de novo* mutations from a single starting genotype (Weilgoss et al. 2013; Long et al. 2015; McDonald et al. 2016; Barghi et al. 2019; Castro et al. 2019), quite unlike the situation in natural populations.

Recent work with *Drosophila* populations has started to clarify the degree of temporal stability of allele frequencies in natural settings. For example, studies on intra-annual changes in allele frequencies have led to interesting observations on cyclical cycles (Bergland et al. 2014; Machado et al. 2021), as suggested earlier by Dobzhansky (1943), although it is difficult to rule out a role for gene flow and spatial microheterogeneity in open populations of flies (Barker et al. 1986; Ives and Band 1986; Coyne and Milstead 1987; Bertram 2021). A recent before/after study of samples of *D. melanogaster* separated by 35 years provides evidence for longer-term changes (Lange et al. 2022), although again there are unresolved issues with respect to selection in very long-term laboratory cultures and on heterogeneity of field sampling sites. Small mesocosms seeded with replicate samples from a synthetic population provide further opportunities for studies of this sort, although the genetic architecture of the base population can be substantially altered with respect to that in the wild (Rudman et al. 2022).

To help close this loop in our understanding of the operational features of natural selection, evolutionary biology could profit from the study of genome-wide selection in natural populations on time scales that minimize ambiguities in interpretations, and in structural settings that do not intentionally introduce novel selection pressures (Kelly 2022; Snead and Alda 2022). For metazoans and land plants, such investigations can require multiple years of sampling and a substantial amount of genome sequencing. Here, we report on nine consecutive years (∼ 35 generations) of population-genomic sequencing of a closed and undisturbed population of *Daphnia pulex,* a cyclically parthenogenetic microcrustacean that inhabits thousands of intermittent ponds across North America. Our results show that although selection is pervasive across the genome, the genome is also in near steady state with respect to selection, with an average selection coefficient experienced by nucleotide sites near 0.0, but with substantial variance in the average selection coefficient across genomic sites and years, and with little temporal correlation in selection within sites. Many aspects of these results are qualitatively consistent with the concept of quasi-neutrality, wherein individual nucleotide sites experience random fluctuations in the direction and magnitude of selection, owing to shifting environmental conditions and/or stochastic changes in patterns of linkage disequilibrium (Kimura 1954; Hartl and Cook 1973).

## Results

### Nine sequential years of population-genomic data

This study utilizes a temporal series of genome-sequence data from annual population samples consisting of 72 to 92 diploid individuals from a single temporary-pond population of *D. pulex,* from Portland Arch, Indiana (Supplemental Text). Each sample was obtained shortly after resting-egg hatch-out (typically mid-March to mid-April) and hence represents first-generation sexually produced offspring emerging prior to the operation of selective events during the subsequent ∼ 3 to 5 generations of clonal reproduction. Horizontal hauls of a zooplankton net throughout the pond minimized any potential effects of microheterogeneity. Generally, the population enters a phase of sexual reproduction and resting egg production by June, as the pond dries up and remains so until the following spring. Genotype frequencies at the time of sampling closely adhere to Hardy-Weinberg expectations, as is true for most other temporary-pond populations of *D. pulex* (Maruki et al. 2022).

After extraction of DNA, the clonal samples were barcoded, multiplexed, and sequenced to an average of 9*×* coverage per clone to generate 100 to 150-bp paired-end reads, which were then mapped to the high-quality reference assembly for a clone taken from an adjacent population (Supplemental Text). Following various filters for quality control (Maruki et al. 2022), allele-frequency estimates were obtained with a maximum-likelihood procedure (Maruki and Lynch 2015).

### Rejecting the hypothesis of random genetic drift

Prior to analyses on the effects of selection on allele-frequency change, we verified that the magnitudes of allele-frequency fluctuations between annual samples are far too large to be consistent with random genetic drift (Supplemental Text). Using a methods-of-moments estimator that eliminates sampling variance as a contributor to the variance of allele-frequency change, we obtained estimates of the *N_e_* necessary to account for the standardized variance of allele-frequency change across all polymorphic sites with adequate coverage. Using only sites for which at least 10 individuals had adequate sample coverage, the upper bound (estimate plus two SEs) to the *N_e_*required to account for the observed allele-frequency changes across all pairs of adjacent years is 500, and increasing the sample-size cutoffs to 20 and 40 individuals leads to estimates of 620 and 540, respectively (Supplemental Table 2). For analyses involving the net change across the first and final sampling years, the upper limits to *N_e_*that can account for the data are 1130, 4420, and 2080, respectively. Each of these genome-wide estimates was based on 2,000,000 to 2,850,000 polymorphic sites.

To determine whether hatching of residual resting eggs from multiple years might influence these estimates (an egg-bank effect), we performed computer simulations with various levels of retention of resting eggs from year to year, comparing the extracted estimates of *N_e_* from the resultant series of allele frequencies with those obtained under the assumption of zero resting-egg retention (Supplemental Text). Egg-bank retention leads to a reduction in the perceived effective population size, as multiple generations of drift occur relative to older recruits. However, the effect is independent of the annual effective size and of the allele frequency, with a fairly small magnitude, *<* 10 and *<* 20% reduction with egg-retention rates as 30 and 50% (which seem highly unlikely).

Thee inferred *N_e_* estimates are inconsistent with standing-levels of variation. Based on levels of silent-site diversity, the PA population is estimated to have a long-term effective population size of ∼ 600, 000, and studies of nine other populations in the same geographic region and with similar ecologies have long-term *N_e_* in the range of 430,000 to 750,000 (Lynch et al. 2020; Maruki et al. 2022). Thus, given the major disparity between the temporal estimates of *N_e_* noted above and those based on standing levels of variation, we conclude that the vast majority of observed temporal changes in allele frequencies reported on below are too large to be reasonably attributable to random genetic drift.

### Mean and temporal variance of the selection intensity

Our goal is to generate a general understanding of the magnitude and degree of selection intensity operating at the nucleotide-site level across the entire genome. Given the time scale of the current study, we cannot expect to estimate with high accuracy the selection coefficient *s* at any site where the latter has an absolute value *<* 0.01 (Lynch and Ho 2020). However, given the > 2, 000, 000 informative sites in the study, it remains possible to estimate the average intensity and variance of selection operating on sites of specific functional significance and in particular chromosomal locations. Nearly all polymorphic sites in the study population contain just two alleles (> 97%; Maruki et al. 2022), but as we cannot yet be certain of the ancestral states at individual sites, the following analyses are oriented on minor alleles at biallelic sites (i.e., those with average frequencies across all years *<* 0.5); the estimated selection coefficients for major alleles have identical absolute values and simply differ in sign.

We emphasize at the outset that selection coefficients estimated in natural populations reflect the net effects of any direct selection operating on the target site and of the aggregated indirect effects of all sites in linkage disequilibrium with the site of interest. With this in mind, we start by considering the distribution of site-specific average annual selection-coefficient estimates (*s*) across the genome. Least-squares regression of logit-transformed minor-allele frequencies yields nearly unbiased estimates of *s* for a site, provided the strength of selection exceeds the power of random genetic drift, and even in the latter case bias towards smaller values is not expected to exceed a factor of two (Lynch and Ho 2020). However, as there is a statistical upper limit to regression estimates of *s* for alleles with very small average minor-allele frequencies (MAFs) in a set of samples, we confined analyses to sites with average MAF > 0.05 and nonzero frequency estimates on at least five sampling dates.

The distributions of site-specific *s* are quite symmetrical, with average values nearly independent of MAF and having an overall mean of *−*0.0025 (SE *<* 0.0001) (Figures 1A,B), i.e., on average, minor-alleles are under weak purifying selection as a consequence of direct effects or (more likely) indirect effects associated with adjacent sites in linkage disequilibrium. Although the widths of the distributions decline with increasing MAF, this is largely a consequence of sampling error, which is inversely proportional to allele frequencies (Supplemental Text). Subtracting the average sampling variance of individual *s̅* estimates from the overall variance provides and taking the square root provides an estimate of the standard deviation of the true distribution of *s̅* among sites, which decline slightly from ∼ 0.045 to 0.030 with increasing MAF (Figure 1B). Thus, the bulk of genomic sites have long-term average net selection coefficients with absolute values *<* 0.05.

**Figure 1.**
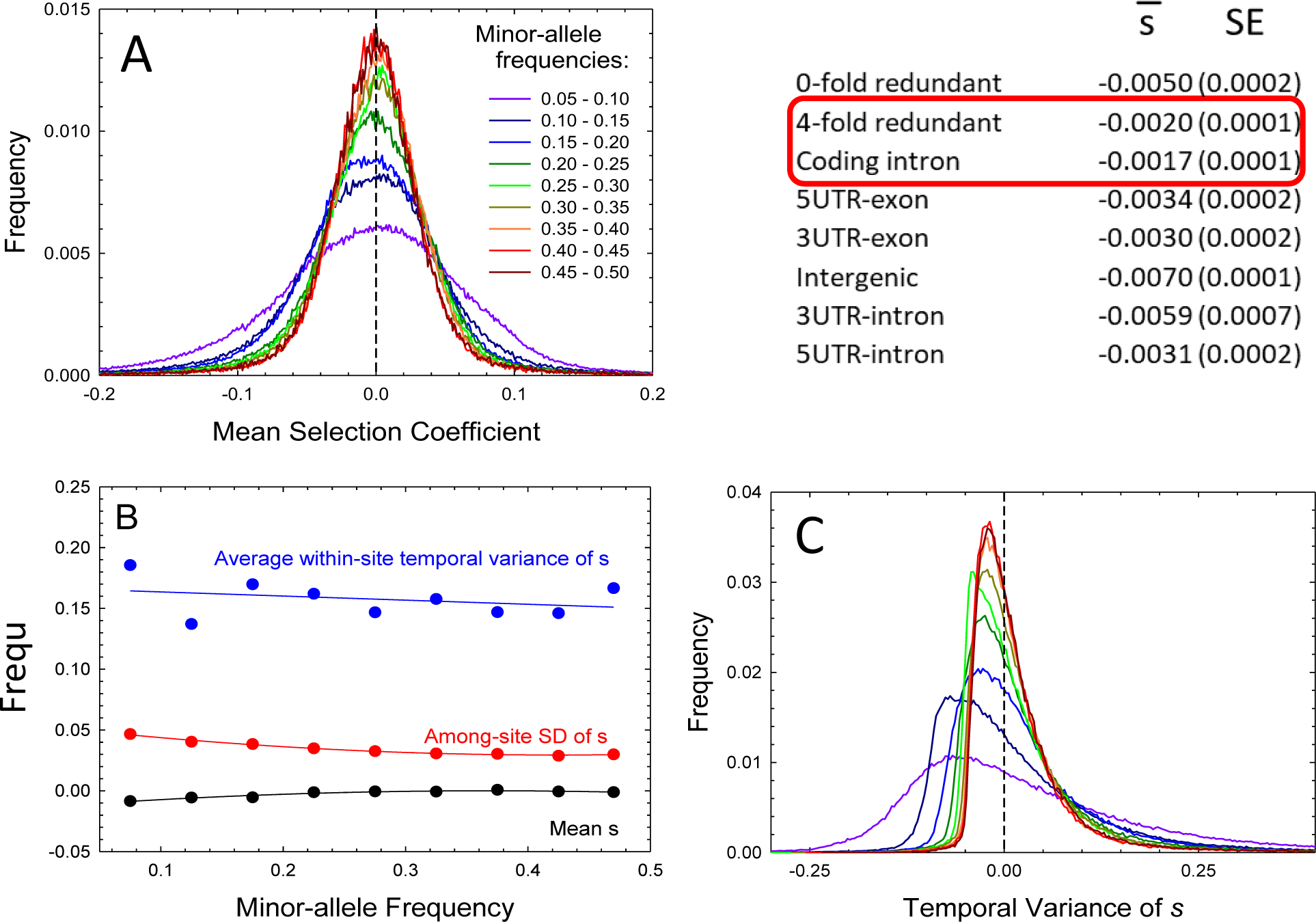
Features of site-specific selection coefficients, across genomic sites. A) Distribution of time-averaged (mean) selection coefficients, *s̅*, for the full set of genomic sites, given for different windows of minor-allele frequencies (MAFs); the table to the right gives averages over all MAFs for different functional categories of sites. B) Means of site-specific *s̅*, the standard deviations of *s* among sites after removal of sampling variance (red points), and the average temporal variance of site-specific *s,* again after removing variance associated with sampling (blue points) as a function of minor-allele frequency. Note that in all cases the standard errors of the estimates are on the order of the width of the plotted points or smaller. C) The distribution of the site-specific estimates of temporal standard deviations of *s* (after removing the contribution from sampling error).

Focusing in on sites with different functional significance (Figure 1, table insert), in all cases, minor alleles experience, in all cases, significant but weak negative selection. This is even true for four-fold redundant sites in coding regions and internal intron sites, which have an overall average *s̅* ≃ −0.002. Average strengths of selection are elevated in all other functional classes (zero-fold redundant sites, UTRs, and intergenic), ranging from *−*0.003 to *−*0.007, i.e., ∼ 1.5*×* to 3.5*×* the means for putatively neutral sites.

By using the eight single-year interval-specific estimates of *s* for each site, it is possible to estimate the within-site temporal variance of selection, 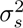, after subtracting out the contribution from variance associated with sampling error. These sample distributions tend to be highly skewed, again with greater variance at lower MAFs resulting from remaining elevated sampling error of the corrected estimates (Figure 1C). Many estimates of 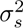 are negative, which is a necessary consequence of the use of an unbiased methods-of-moments estimator with an expectation near zero. However, the average values of *σ_s_* across sites are nearly independent of MAF, with a mean value ≃ 0.16. These results are qualitatively similar to, although much more precise than, those obtained 35 years ago from temporal series of allozyme-allele frequencies in different *Daphnia* populations (Lynch 1987).

The fact that both the means and standard deviations of *s̅* are nearly independent of MAF is consistent with selection operating on linkage blocks of sites containing a range of allele frequencies, all subject to the same net selective effects associated with their cumulative features. The average allele is then subject to a long-term mean absolute selection coefficient of ∼ 0.0025, with substantial variance of *s̅* among sites (standard deviation ≃ 0.04), and a larger average between-year, within-site standard deviation of *s ≃* 0.16. Note that all of these selection coefficients are measured on an annual time scale, so with ∼ 4 generations per year, *s̅* and *σ_s_* should be reduced by this same factor to yield per-generation estimates, which means that the average absolute strength of selection operating on individual genomic sites is *<* 0.001 per generation. The potential effects of a residual egg bank are presented in the Supplemental Material, where it is shown that these are minimal (*<* 10% upward bias) for *σ_s_,* and although more pronounced for *s̅* estimates, not enough to alter the conclusions noted above.

### Temporal covariance of selection intensity

In principle, selection coefficients at individual nucleotide sites can exhibit temporal correlations owing to persistent environmental and/or linkage effects. To determine whether the annual selection coefficients associated with individual sites covary from year to year, we estimated the within-site covariance of *s* separated by intervals of *T* = 1, 2, 3, and 4 years, using unbiased estimators (Supplemental Text). A linear representation of the selection intensity operating on a site between adjacent time points *i* and *i* + 1 is

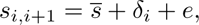

where *s̅* is the mean selection coefficient over time (the subject of the previous section), *δ_i_* is the deviation from the expectation in generation *i* associated with environmental and/or background genetic effects on selection, and *e* is the sampling error associated with estimation. The expected within-site covariance of annual selection coefficients separated by intervals of *T* years is *σ*(*δ*_0_*, δ*_1+*T*_). That is, the temporal covariance in selection coefficients for a particular site is equal to the average covariance of deviations of observed interval-specific estimates of *s* from *s*, using the full set of pairs of estimates 1 + *T* sampling points apart across the entire time sequence. (Note that *T* = 1 means that pairs of selection coefficients are separated by an entire annual interval, e.g., *s* based on years *x* and *x* + 1 vs. *s* based on years *x* + 2 and *x* + 3; this avoids the use of overlapping years in adjacent time intervals, which can lead to substantial bias from negative sampling covariance of estimates of *s*).

Relative to the temporal variance in *s*, which as noted above is close to 0.025, temporal covariances are much smaller and again essentially independent of the MAF, with mean values of -0.0026 (SE = 0.0005), 0.0056 (0.0008), 0.0041 (0.0006), and -0.0039 (0.0006) for *T* = 1, 2, 3, and 4, respectively (Figure 2). Again, it can be seen that the distributions of the temporal covariance estimates are quite symmetrical, with the widths of the distributions declining with increasing MAF. Such a decline is in part a consequence of the reduction in sampling variance, but may also be due to the fact the bulk of high-frequency MAFs in *D. pulex* populations have nearly neutral fitness effects (Lynch et al. 2017), as implied by the behavior of *s̅* in Figure 1B. Notably, there is no gradual reduction in the temporal covariance of *s* with time, as might be expected if correlations between environmental factors consistently dissipate over time. Moreover, two of the average covariances are negative, suggesting that for this particular temporal sequence of nine samples, environmental factors influencing this population on time scales separating *s* by 2 and 5 years yielded negative correlations in selective forces (on average), whereas those separated by 3 and 4 years tended to have positive effects. These results do not rule out the possibility of selection covariance between adjacent generations within-years, which this study is unable to ascertain, although prior work with electrophoretic variants suggests that this is not the case (Lynch 1987).

**Figure 2.**
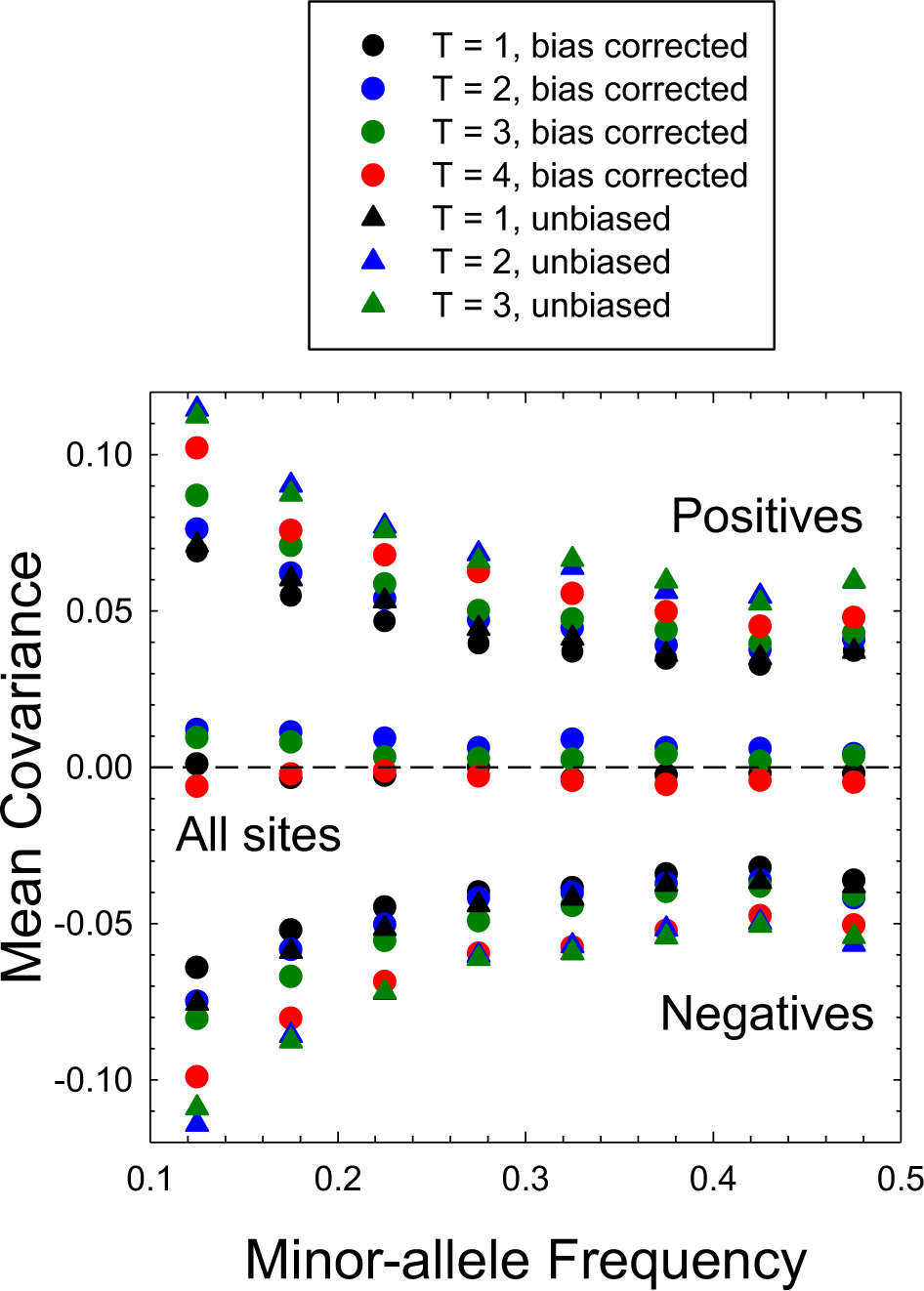
Average temporal covariance of selection coefficients for individual sites over four different time intervals (*T* = 1,2,3,4). Results are given for the averages over all sites, and separately for the averages conditional on being positive or negative. The range narrows to the right owing to the reduction in sampling variance when minor-allele frequencies are high. The dashed line is the reference for a covariance of zero. In all cases, the standard errors of the estimates arc on the order of the width of the plotted points or smaller.

Finally, it should be noted that a negative selection covariance does not necessarily imply a change in the sign of *s* across generations, as previously inferred (Buffalo and Coop 2020), but can also arise when the sign of *s* remains the same but with more extreme values of *s* in one interval being followed by less extreme values in the next. This can be readily seen from the relative incidences of +*/*+, *−/−*, and opposite-sign pairs for the full sets of sites exhibiting positive and negative selection covariances across all time intervals. Independent of the MAF, for the positive covariances, 31.5% of paired *s* are +*/*+, 31.5% are *−/−,* and 37.0% are +*/−*. For the negative covariances, these numbers are 18.8, 18.2, and 63.0%, respectively (all SEs are *<* 0.1%). Thus, although sequences of +*/−* pairs predominate for sites with negative covariances, they are also most common for sites with positive covariance.

### Genome-wide temporal covariance of selection

To gain an appreciation of the extent to which this population is experiencing consistent forms of selection across time, we also applied the method of Buffalo and Coop (2019) to estimate the genome-wide covariance of allele-frequency changes across time intervals. (A similar approach was introduced by Bertram (2021)). The approach taken is complementary to that in the preceding section, but rather than focusing on individual nucleotide sites, the goal here is to evaluate whether the changes in frequencies for the full set of polymorphic sites are correlated across two sets of time intervals, as might be expected for a polygenic trait experiencing directional selection. With allele-frequency changes available for eight different single-year time intervals, 28 pairwise comparisons could be made, ranging from seven involving single generation intervals to one separated by seven intervals, with an average number of 2.5 *×* 10^6^ sites used per individual covariance estimate. Using the the approaches advocated by Buffalo and Coop (2020), we performed such analyses for the pooled set of all genomic sites as well as for those in narrower MAF windows, in each case standardizing the covariance measures by dividing by *p*(1 *− p*), as the magnitude of selection is expected to be proportional to the heterozygosity (Supplemental Text).

The majority of such standardized covariance estimates are either very close to zero or negative (Figure 3). For time intervals of one and two years, estimates for the pooled set of sites are significantly less than zero, and the estimates for all MAF classes are negative as well. For all longer intervals, however, none of the genome-wide estimates are significantly different from zero, and the estimates for the individual MAF classes are distributed around zero as well, with most being not significantly different from zero. The overall conclusion here is that the study population reveals no evidence of consistent directional selection at the polygenic level, little temporal correlation beyond a two-year interval, and with the negative covariances for short intervals being potentially consistent with some form of fluctuating stabilizing selection.

**Figure 3.**
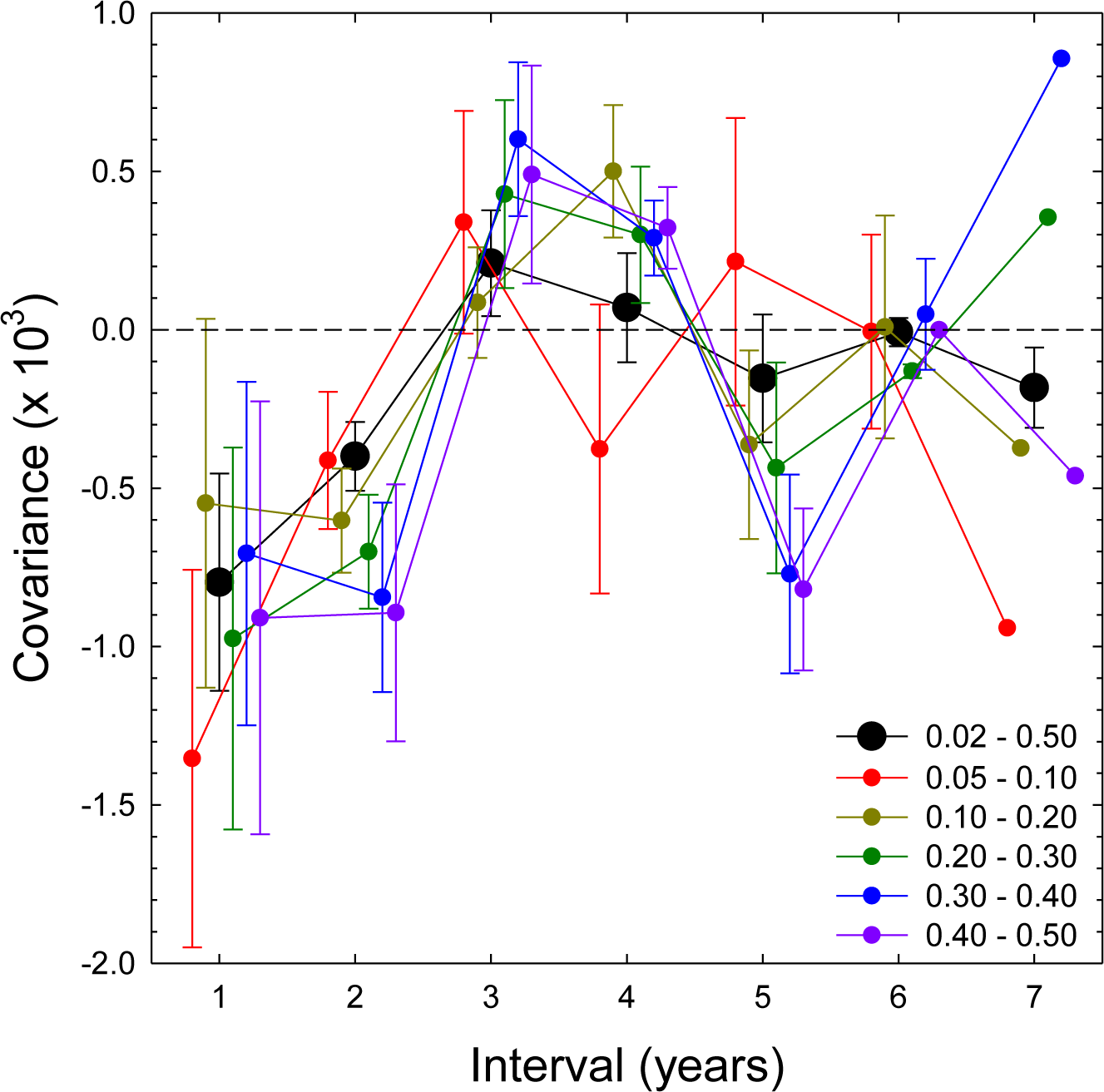
Genome-wide temporal covariance of allele-frequency changes across intervals with increasing length. With eight consecutive annual sampling points, there are seven ways to compute single-year changes in adjacent years, six ways to compare single-year changes separated by two years, etc., and the plotted values denote the means and SEs of such sets. (There is only one way to compare changes across a seven year interval, the change between the first two years vs. that between the final two years, and so the SE is only reported for the full-genome analysis, based on the SE of the means of the window-specific estimates). Results are given for the full set of polymorphic sites as well as for narrower windows of minor-allele frequencies (denoted in the inset). For each time interval, the results for the different MAF windows are offset slightly on the x axis for visualization purposes.

### Chromosome-wide distributions of selection intensity

To evaluate the degree to which the average strength of selection operating on nucleotide sites is a function of that operating on linked sites, we estimated the average absolute value of *s* over all sites within blocks of 100 consecutive sites with MAFs > 0.1, proceeding along the full lengths of chromosomes with adjacent blocks overlapping by 50%. We also drew 10^7^ random sets of such SNPs from the entire genomic pool of data to calculate the distribution of average *s* in the absence of linkage effects. By comparing the observed (|*s̅*| - 2 SEs) for each window (i.e., making the estimate less extreme) with the cumulative random probability distribution, we were then able to determine the conservative probability *P* of obtaining a window-specific *s̅* more extreme than that observed. The overall survey evaluated the features of 27,103 windows with average and standard deviation of length 6.84 (SE = 0.05) and 8.26 kb. Such windows typically will contain 0 to 3 protein-coding genes, as the average gene span in *D. pulex* is ∼ 10 kb. After correcting for multiple comparisons, we used a critical *P* value of 1.85 *×* 10*^−^*^6^ as a cutoff for windows with *s* significantly elevated over the random expectation.

This analysis revealed both large linkage blocks and punctate regions of one or more adjacent windows exhibiting significantly strong collective selective effects (Figure 4A). Correlation analysis of nonoverlapping windows indicates a clear spatial clustering of *s,* extending out to a length scale ≃ 20 windows, equivalent to ∼ 137 kb (Figure 4B). Regions of significant selection are found on every chromosome arm, and centromeric regions often encompass multiple patches of significant spans of selection.

**Figure 4.**
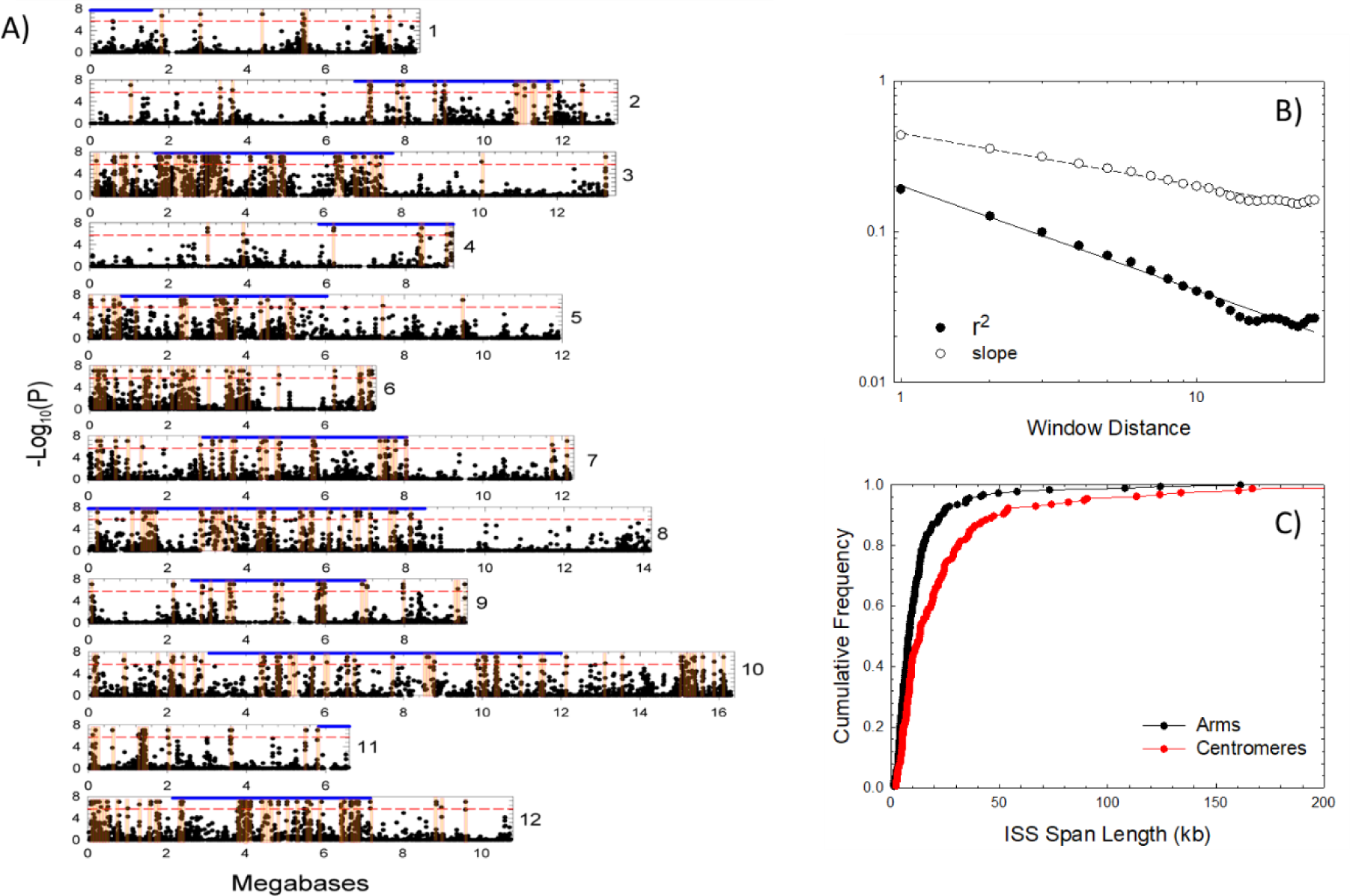
Chromosome-wide scans of window-specific intensities of selection. **A)** For the set of SNPs within each window, the average estimate of absolute values of selection coefficients was compared with the probability distribution based on samples of randomized sites, with *P* denoting the probability of achieving the observed measure by chance. The dashed red lines denote the critical cutoff points for significance after correcting for multiple comparisons. Horizontal blue bars denote the approximate locations of centromeric regions based on multiple laboratory crosses (Molinier et al. 2021). **B)** Decline of the correlation of window-specific selection strengths with increasing distance, reported as both the squared correlation coefficients and the slopes of the regressions of pairs of windows at the window distance designated on the x-axis, where *x* = 1 denotes adjacent (nonoverlapping) windows. **C**) Cumulative frequency distributions for the lengths of ISS (islands of strong selection) blocks residing on chromosome arms vs. windows.

To gain further insight into the spatial patterning of selection and its consequences, we delineated islands of strong selection (ISSs) as chromosomal blocks harboring adjacent windows with high significance, with each span constructed by starting with windows deemed individually significant and then extending the span to adjacent segments with *P <* 10*^−^*^4^. This treatment with a somewhat reduced *P* level in adjacent windows is necessary because: 1) the breakpoints associated with islands of interest will often be embedded within end windows, thereby diminishing their overall probability levels; and 2) such a borderline window may be immediately followed by another string of significant windows and so may be a false negative (even though it is still significant at this lower probability level). Although the acceptance criterion used here is somewhat arbitrary, the approach used is quite conservative, as the multiple-comparison correction conditional on a small number of local windows is much less stringent than that for the entire genome.

This sliding-window analysis led to the identification of 341 ISSs, 145 of which reside on chromosome arms, ranging from 1.4 to 41.5 kb in length. The average number of ISSs per Mb within centromeric regions is 1.7*×* greater than that on chromosome arms, 3.0 (SE=0.7) vs. 1.8 (0.4), and the average ISS span within centromeric regions is more than twice that for ISSs on chromosome arms: 24.2 kb (SE = 2.8) vs. 9.6 (0.6) (Figure 4C). As a result, a greater fraction of centromeric regions is occupied by ISSs than is the case for chromosome arms, 0.073 (0.016) vs. 0.018 (0.004). This may in part be a consequence of the lower levels of recombination in centromeric regions, and the resultant effects of linked selection. Although little crossing over is revealed in centromeric regions in single-generation laboratory crosses, the long-term average rate of recombination is suppressed ∼ 2-fold relative to the average rate on chromosome arms (Lynch et al. 2022).

### Polymorphism and divergence of potential targets of selection

There are several unique features of the genes contained within ISSs (Table 1). First, ∼ 50% of them are young gene duplicates (*<* 2% sequence divergence), in contrast to only ∼ 10% of such genes in non-ISS regions. A large fraction (0.46) of these ISS duplicates have no obvious orthologs in other metazoans, and may be *Daphnia*-specific genes. (To search for orthologs, we subjected each annotated *D. pulex* gene to Blast analysis against the non-redundant NCBI protein-coding database, regarding any gene with *<* 30% sequence coverage (with E-value > 0.00001) for all non-*Daphnia* species as potentially unique to *Daphnia*). Second, ISS genes exhibit significantly lower levels of average silent-site diversity (*π_S_*) but significantly higher levels of replacement-site diversity (*π_N_*) than non-ISS genes, with the *π_N_ /π_S_* ratio in the former being 1.2*×* (arms) to 1.9*×* (centromeres) higher than in the latter. Third, with respect to average divergence from the closely related outgroup species *D. obtusa,* the ISS genes exhibit higher levels of *d_N_, d_S_,* and *d_N_ /d_S_*, the latter being 1.3*×* (arms) to 2.0*×* (centromeres) higher than than in non-ISS regions. As a consequence of the parallel elevation of both *π_N_ /π_S_* and *d_N_ /d_S_*, the neutrality index, NI = (*π_N_ /π_S_*)*/*(*d_N_ /d_S_*) (with NI = 1 implying neutrality, on average) is similar between ISS and non-ISS genes. As levels of polymorphism and divergence are cumulative products of evolutionary factors operating over long time scales, the existence of such distinguishing features of ISSs suggests that the regions involved are not simply fortuitous responders to the unique environmental conditions encountered during the limited eight-year span of this study, but instead reflect longer-term consequences of ecological and/or linkage-disequilibrium issues.

**Table 1.**
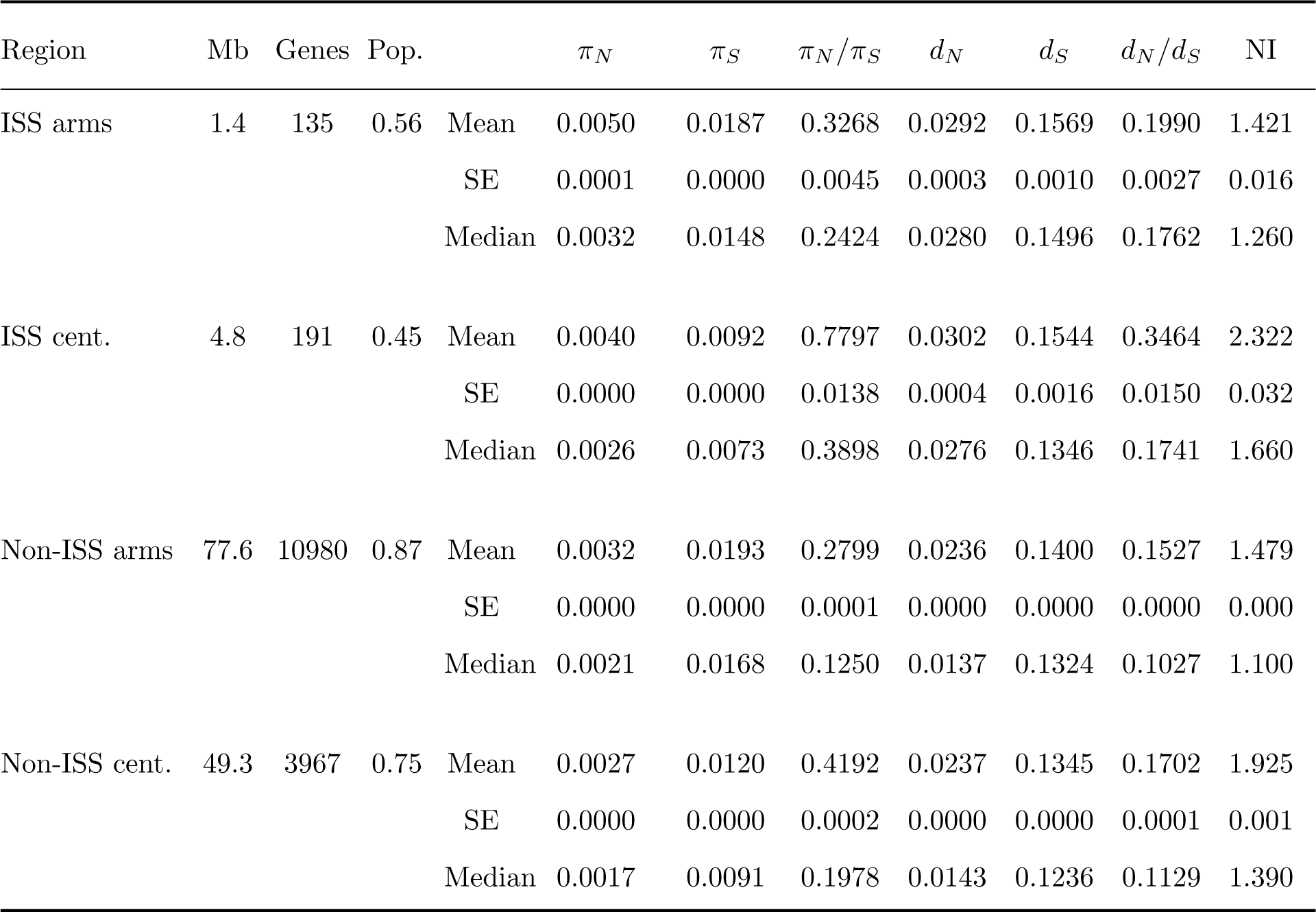
Estimates of within-population diversity measures (*π*) for synonymous (S) and non-synonymous (N) sites in protein-coding genes, as well as for divergence (*d*) statistics from comparisons with orthologous genes in the outgroup species *Daphnia obtusa*. Total length of regions is in units of Mb; Genes refers to the total number of genes analyzed; Pop. refers to the fraction of genes with uniquely mapped reads from the population surveys; NI denotes the mean neutrality index (ratio of *π_N_ /π_S_* to *d_N_ /d_S_*). Data are given separately for chromosomal arms vs. centromeres, and for genes in islands of significant selection (ISSs) vs. non-ISSs. All statistics are averages over all genes, with SEs of the means given below the means.

The preceding observations are qualitatively consistent with predictions from theoretical population genetics, while also motivating the need for further work in this area. For example, for the special case in which mutation is unbiased (with rate *u* per nucleotide site) and *s* = 0, the steady-state distribution of allele frequency *x* with drift, reversible mutation, and fluctuating selection is given by

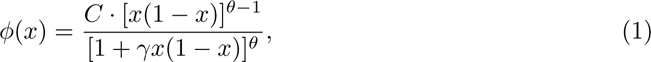

where 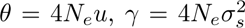, and *C* is a normalization constant such that the integration of Equation (1) over the full allele-frequency spectrum is equal to 1.0 (Karlin and Levikson 1974; Takahata and Kimura 1979). Multiplying *φ*(*x*) by 2*x*(1 *− x*) and integrating over *x* = 0 to 1 yields an estimate of the expected nucleotide diversity (*π*) under fluctuating selection.

The average level of silent-site diversity across the genome in the study population is 0.014, and under the assumptions of neutrality, this measure is traditionally interpreted as being an estimate of *θ,* which after factoring out the known mutation rate (Keith et al. 2016) yields *N_e_ ≃* 600, 000. If this conventional interpretation of *π_s_* is close to correct, the above estimates of *σ*^2^ imply that *γ* may often be on the order of 10^4^ or even higher. How might such variation in selection intensity influence standing-levels of diversity? From Equation (1), it can be seen that if 0 *< γ <* 1, the denominator will generally be close to 1, and *φ*(*x*) will be closely approximated by the numerator alone, which is equivalent to the neutral expectation. Solution of Equation (1) shows that for *γ* = 10^1^, 10^3^, 10^5^, and 10^7^, expected levels of heterozygosity decline to 0.99*θ*, 0.94*θ*, 0.89*θ*, and 0.84*θ*, respectively.

Although this analysis suggests that fluctuating selection causes standing levels of silent-site diversity to underestimate *θ* (and hence the *N_e_* estimate) in the study population by as much as 20%, the actual depression is likely greater. One issue with Equation (1) is that it presupposes some extrinsic measure of *N_e_* in the definition of *γ,* whereas the latter is actually defined in part by fluctuations in selection, i.e., by 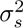 itself. This can be shown with computer simulations in a Wright-Fisher framework with consecutive episodes of reversible mutation, selection (assuming a Gaussian distribution of *s*), and drift (Supplemental Text). By following the behavior of a selected site and a completely linked neutral site (mutating independently at the same rate), the long-term mean values of *π_S_* and *π_N_* can be evaluated.

In the absence of selective interference, the mean nucleotide diversity at the silent site is 4*Nu/*(1 + 4*Nu*), where *N* is the effective population size in the absence of selection. However, the simulation results show that with increasing *N*, mean silent-site diversity begins to decline once *N* exceeds ∼ 1000/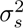, with a 100-fold increase in *N* beyond this point leading to a nearly ∼ 100-fold decline in *π_s_* (Figure 5). Below this critical point, the behavior of diversity at the linked neutral and selected sites remains equivalent, so that *π_N_ /π_S_* = 1, but with increasing *N*, *π_N_ /π_S_* begins to exceed 1, and with large (but realistic) values of 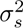, *π_N_ /π_S_* can exceed 10. In this domain, *π_N_* is depressed below the neutral expectation of 4*Nu*, as anticipated by Equation (1), but still grows slowly with increasing *N* while *π_S_* is declining. Although nonzero *s* results in lower *π_N_ /π_S_*, the tendency for *π_N_ /π_S_* to exceed 1.0 still occurs with large *N*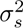. This type of behavior results because while fluctuating selection tends to encourage the maintenance of variation at selected sites by imposing a sort of frequency-dependent selection, it also magnifies the fluctuations in allele frequencies, which reduces the depth of gene genealogies, thereby reducing the expected value of *π_S_* (Taylor 2013).

**Figure 5.**
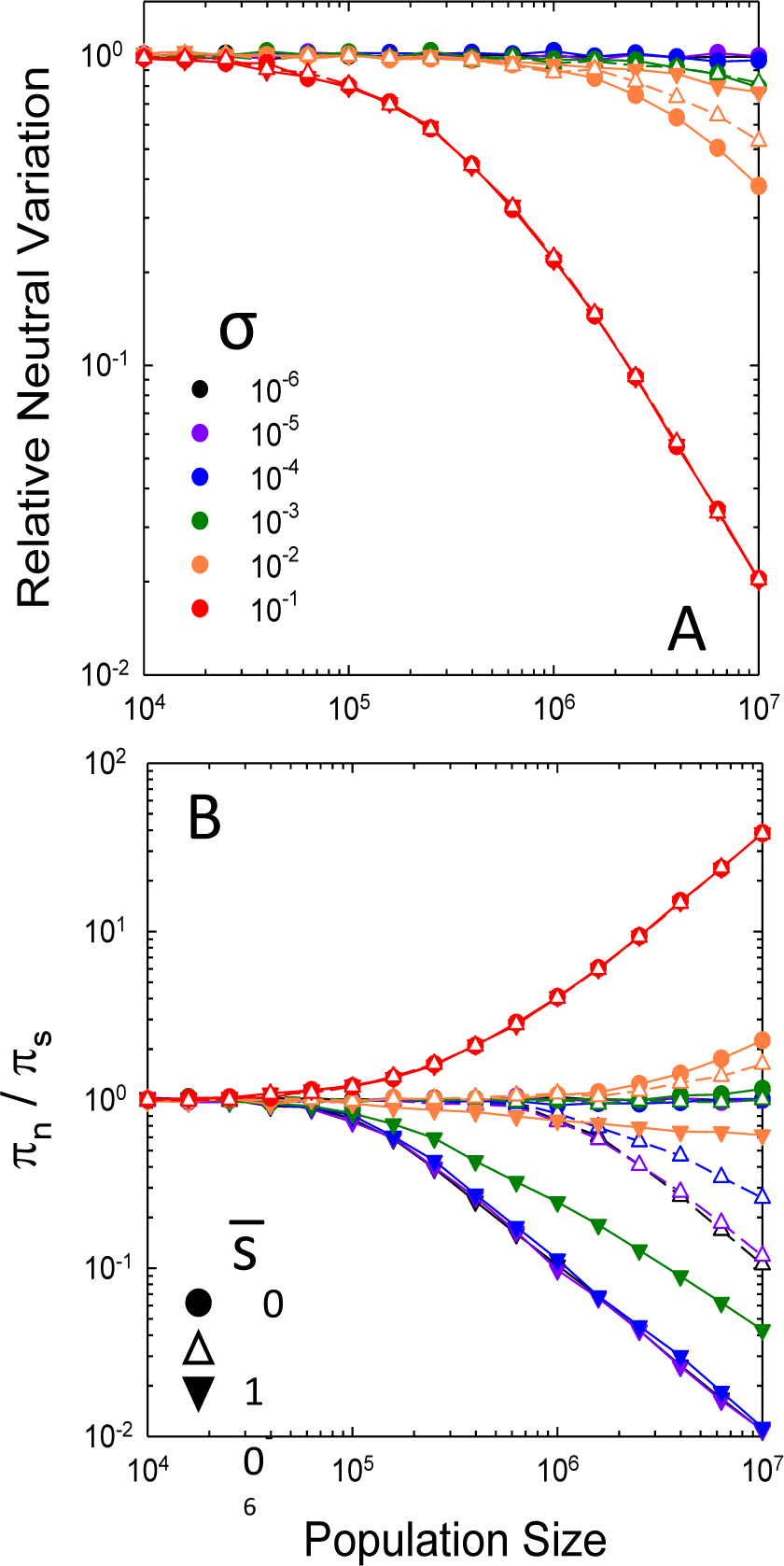
Response of expected within-population measures of nucleotide diversity as a function of the temporal standard deviation of the selection coefficient (σ_*s*_) and the population size (*N*). **A)** Nucleotide diversity at a linked silent site (π_*S*_) relative to the neutral expectation at drift-mutation equilibrium under free recombination. **B)** Ratio of diversity at the selected site to that at a completely linked neutral site, π*N*/π*S*. Color legend in the upper panel designates various levels of the temporal standard deviation of the selection coefficient *s*; inset in the lower panel designates the symbol shapes used for three levels of the average selection coefficient *s̅*.

These predictions from theory are concordant with the observations on nucleotide diversity in Table 1 – a decline in *π_S_*, an increase in *π_N_*, and a resultant dramatic increase in the ratio of the two in ISS vs. non-ISS regions. Theory also predicts that fixation probabilities will increase with fluctuating selection (Kimura 1954; Karlin and Levikson 1974; Takahata et al. 1975; Dean et al. 2018; Cvijovć et al. 2015), which is consistent with our observations on *d_N_*. There has been some suggestion that fluctuating selection will decrease the ratio of polymorphism to divergence, thereby leading to the false impression of positive selection (Huerta-Sanchez et al. 2008). However, the latter work does not disentangle the effects of fluctuating selection on *N_e_*, and the predicted pattern is not apparent in the data herein.

## Discussion

Theoreticians have long appreciated the potential effects of fluctuating selection on patterns of nucleotide diversity and divergence, although in the absence of empirical data on the magnitude of the variance in selection intensity (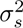), most studies in molecular population genetics assume that fluctuating selection is of negligible significance. This is a concern because the movement of allele frequencies by both random genetic drift and selection is proportional to the heterozygosity at a nucleotide site, and for the special case of *s̅* = 0, the influence of fluctuating selection can be nearly indistinguishable from the effects of drift associated with gamete sampling (Tier 1982). Despite this conceptual problem, hundreds of studies have interpreted patterns of population-genetic variation in terms of drift, mutation, and deterministic selection. For example, such a strategy remains central to virtually all studies using measures of nucleotide variation/covariation at putatively neutral sites to estimate *N_e_* after factoring out known estimates of mutation and/or recombination rates. It is also central to studies relying on estimates of *π_N_, π_S_, d_N_*, and *d_S_,* and their various ratios to derive inferences about the relative strength of positive and purifying selection within and among genes and species.

The current study presents one of the first genome-wide analyses of temporal variation in selection in an isolated and unmanipulated natural population, and in doing so illustrates the significant ways in which this under-appreciated issue can influence the interpretation of parameter estimates in molecular population genetics. Shorter-term studies in the fruit fly *Drosophila melanogaster* have recently appeared, although there are uncertainties with respect to gene flow and microhabitat heterogeneity (Bergland et al. 2014; Machado et al. 2021) and the reliance on artificially constructed base populations (Rudman et al. (2022). Our results show that selection coefficients operating at the nucleotide-site level in *D. pulex* are quite small in terms of absolute values, with the average annual selection coefficient operating on minor alleles often being in the range of *−*10*^−^*^4^ to *−*10*^−^*^3^ or smaller (and ∼ 25% of that on a per generation basis). For sites that are most likely to be functionally neutral, estimates of *s* are closer to but still significantly different from 0.0 on average, presumably reflecting the effects of linkage disequilibrium with functionally relevant sites. However, given the long-term effective population size of *D. pulex,* selection coefficients as small as 10*^−^*^4^ represent strengths of selection well beyond the power of random genetic drift.

Although we report on the results of just a single population, there is no evidence that the results are peculiar consequences of the cyclical nature of the sexual life-cycle phase in *Daphnia*. For example, despite the brief bouts of asexuality each generation, the population-genetic features of *D. pulex* are quite similar to those of the well-studied *Drosophila melanogaster,* which on an annual basis actually has significantly less per-generation recombination per nucleotide site than the study population (Lynch et al. 2017; Maruki et al. 2022). Nor does the possible (albeit unlikely) existence of a moderately large resting-egg bank alter our results in a substantive way (Supplemental Text). Notably, in analyses of the genome-wide temporal covariance of allele-frequency change in a natural population of *D. melanogaster* (Bergland et al. 2014), Buffalo and Coop (2020) and Bertran (2021) found a similar pattern to that reported in Figure 3 – on average, weak but negative covariance over the shortest interval and near-zero covariance thereafter. Likewise, although the methods employed were quite different, a ten-year genomic survey of frequency changes for common alleles in the monkey flower (*Mimulus guttatus*) revealed substantial fluctuating selection associated with numerous islands of linked selection and modest temporal covariance of *s* (Kelly 2022)

Thus, observations from at least three species are consistent with the hypothesis of significant fluctuations in selection intensities operating at the level of nucleotide sites, often with very little autocorrelation in selection across years. Nonetheless, there is a compelling need for additional empirical work with different kinds of organisms to determine whether general phylogenetic patterns exist in genome-wide distributions of key composite parameters such as 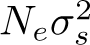.

Taken together, our results appear to be quite compatible with the scenario of quasi-neutrality envisioned by Wright (1948) and Kimura (1954), whereby allelic variants have temporal average selection coefficients close to zero, while experiencing significant selection pressures in some generations. Although there is some temporal covariance of selection experienced by individual nucleotide sites in *D. pulex,* this is on average quite small relative to the temporal variance of *s*, and might be revealed to be even closer to zero with a longer temporal series of data, rendering the overall temporal pattern of selection close to the idealized model of Wright and Kimura.

What seems clear, however, is that rather than being entirely a consequence of simple stochastic effects of direct selection operating on individual nucleotide sites in isolation, as assumed in prior models, the observed features of each site are collectively driven to a large extent by the influences of selection operating on neighboring sites under linkage disequilibrium. Linkage blocks of sites under significant selection over the 9-year sampling period were often on the scale of 50 kb or smaller, although some spans extend up to 0.5 Mb, especially in centromeric regions. Such observations are consistent with prior work in *D. pulex,* including in the study population, indicating that significant linkage disequilibrium is quite strong at distances *<* 10^3^ bp and then gradually dissipates out to ∼ 250, 000 bp (Lynch et al. 2022).

Given that 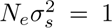 is the approximate benchmark above which fluctuating selection begins to play a role in determining patterns of molecular evolution, our observation that the magnitude of 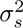 is commonly orders of magnitude greater than the power of random genetic drift, 1*/*(2*N_e_*) suggests that such fluctuating selection may play just as substantial a role as the effects of finite population size in determining the fates of mutant alleles. Of particular concern is the demonstration that increasing levels of fluctuating selection can differentially influence *π_S_* and *π_N_* in ways that can lead to the false impression of relaxed selection (e.g., *π_N_ /π_S_*approaching 1.0) or strong balancing selection (*π_N_ /π_S_ >* 1.0). These observations motivate the need for further theoretical investigation into the extent to which fluctuating selection alters the ways in which patterns of molecular variation within and between species should be interpreted. The challenges are considerable, given that most existing theory treats *N_e_* as a fixed parameter, ignoring the influence of 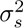 on *N_e_* and the additional complications associated with non-zero *s̅* and non-Gaussian and episodic distributions of *s.* Because virtually all population-genetic theory in this area focuses on single loci, there is a real need for the development of models that incorporate the effects of linked sites jointly influenced by fluctuating selection. Virtually no nucleotide site can be permanently immune to such effects.

Our results are also relevant to the common use of “Manhattan plots” based on chromosome-wide scans in studies of isolated populations and/or species to identify candidate regions experiencing strong positive selection. As the ISSs observed in this study are the outcomes of fewer than 40 generations of evolution, this raises the possibility that many metapopulation studies may not be identifying long-term targets of local adaptation, but simply identifying the serendipitous outcomes of the most recent generations experienced by the populations involved. Thus, in the future, it will be useful to evaluate whether chromosomal regions that are most subject to selection within a population over short time scales (as observed here) are correlated with islands of divergence among subpopulations.

Finally, although science increasingly relies on rapid experimental analyses, nature proceeds at its own pace. Our results strongly suggest that multi-generational population-genomic surveys, combined with ecological observations, will be essential to understanding the population-genetic processes and adaptations that are associated with stable and/or changing environments. Indeed, substantially longer surveys than that reported on here will likely be required to determine how and why the chromosomal targets of selection shift over ecologically relevant time scales. Given the current support of long-term ecological observatory networks (e.g., NEON in the United States) and the much more substantial investment in understanding the history of the universe (e.g., the Webb telescope), the additional costs of gaining insights into key microevolutionary processes in natural populations are not unreasonable. *Daphnia* populations inhabiting large permanent lakes offer considerable opportunities here, as their resting eggs can remain preserved in undisturbed sediments for up to 200 years. With enough sampling from sediment cores with stable stratigraphies, population-level studies might be extended over time periods exceeding several hundreds of generations (Roy Chowdhury et al. 2015; Yousey et al. 2018; Chatuvedi et al. 2021; Isanti-Navarro et al. 2021; Wersebe et al. 2023).

## Materials and Methods

This study relies on a temporal series of nine samples from an isolated temporary pond population of *D. pulex,* derived from newly arisen annual cohorts of resting-egg hatchlings, in the Portland Arch Nature Conservancy. Full genomic sequences were obtained for ∼ 90 individuals from each sample, using methods more fully described in the Supplemental Text but following the same protocols as in prior work with this and other related populations (e.g., Lynch et al. 2017; Maruki et al. 2022). Methods for estimating selection coefficients and their sampling variance are derived from Lynch (1987) and Lynch and Ho (2020), using modifications outlined in the Supplemental Text to reduce sampling bias and minimize sampling error.

## Acknowledgments.

We thank Cora Anderson, Bret Coggins, David Molik, and Ken Spitze, for assistance with sample collection, Matthew Carmody, Matthew Donahue, Nhu-Y Nguyen, Rebecca Shelley, and Emily Williams for help with clone maintenance. Jeff Jensen and Jay Taylor provided helpful comments on the manuscript. DNA Sequencing libraries were prepared by the Notre Dame Genomics and Bioinformatics Core Facility. The work was supported by National Institutes of Health grants R01-GM101672 and R35-GM122566-01 and National Science Foundation grant IOS-1922914.

