## Supplementary File for "The Genome-wide Signature of Short-term Temporal Selection"

### SUPPLEMENTAL TEXT

#### Population Sampling, Sequencing, and Allele-frequency Analysis

Population samples were collected annually by monitoring the until resting-egg hatchlings had emerged in large enough numbers to be obtained after large numbers of hauls of a zooplankton net throughout the pond; based on their small size, all of the individuals sampled were almost certainly first-generation individuals. The clones were then isolated into individual beakers in the lab and grown to high enough densities for extraction and library preparation of DNA using a Bioo or Nextera kit, followed by tagging with unique oligomer barcodes. Pooled sequencing then involved paired-end short (100 or 150 bp) reads obtained using an Illumina NextSeq, HighSeq, NovaSeq, or DNBseq platform at either the Center for Genomics and Bioinformatics at Indiana University (2013-2017), the Beijing Genomics Institution (2018-2020), or Tgen (2021). Samples have been sequenced on Illumina NextSeq, HiSeq, NovaSeq or DNBseq platform. The sequencing files are available at NCBI under the accession number PRJNA684968.

We removed residual adapters and trimmed low quality sequences as described in Maruki et al. (2022). All cleaned reads, including paired and unpaired reads, were then mapped to the reference genome of *D. pulex* KAP4 (available at NCBI under accession GCF 021134715.1; a high-quality genome assembly at near-complete chromosome level) with BWA (Li and Durbin 2009). We merged and sorted alignments using Picard (https://broadinstitute.github.io/picard/), and PCR duplicates were removed with default parameters using Picard. RealignerTargetCreator and IndelRealigner of GATK (McKenna et al. 2012) was used to realigned filtered alignments. In addition, we clipped overlapping alignments by applying BamUtil (Jun et al. 2015). We generated mpileup files with Samtools (Danecek et al. 2021). We called candidate alleles using MAPGD and GFE (Maruki et al. 2015).

Although we attempted to sequence $\sim 96$ clones of each year, a few clones were removed in the following analysis. 1) The clones with low depths of sequence coverage ($< 3\times$) and/or a sum of the goodness-of-fit values across the genome $< -0.4$ (implying nonbinomially sampled allele frequencies within clones, and potential contamination) were discarded. 2) We estimated the pairwise relatedness of clones using the "relatedness function of MAPGD (Ackerman et al. 2017), and removed a single member (with lowest sequence coverage) of clone pairs with relatedness estimates $> 0.125$ to eliminate the possibility of sampling half- or full-sib hatchlings. We estimated the pairwise relatedness of clones using relatedness of MAPGD; only the clone with the highest coverage in any cluster with relatedness estimate $> 0.125$ was retained. 3) We also screened for the possibility of obligately asexual clones, using a pool of 5,263 unambiguous markers for such clones (Xu et al. 2015), but found no such clones. Finally, to further guard against potential mapping issues, we focused only on sites in the 15 to 85 percentile range of sequencing coverage within each sample. Of the 848 clones sent out for sequencing, 1.5% were lost to inadequate coverage, 0.5% were lost to potential contamination, and 6.6% due to relatedness (61% of which occurred in samples in the first two years of study, when samples were gathered very early).

For estimates of nucleotide divergence, we used *Daphnia obtusa* as an outgroup, as it is $\sim$ 12% divergent from *D. pulex* at silent sites. The FASTQ files for the *D. obtusa* sequence reads are available at the NCBI SRA (accession number SAMN12816670), and the complete genome assembly is available at the NCBI GenBank (accession number JAACYE000000000).

**Supplemental Table 1.** Annual numbers of sequenced diploid genotypes and average depth of sequencing coverage per mapped nucleotide site.

| Year | Number of Clones | Depth of Coverage |
|---|---|---|
| 2013 | 78 | 9.7 |
| 2014 | 72 | 9.8 |
| 2015 | 92 | 7.7 |
| 2016 | 87 | 8.1 |
| 2017 | 90 | 6.7 |
| 2018 | 81 | 10.9 |
| 2019 | 88 | 6.9 |
| 2020 | 94 | 9.7 |
| 2021 | 93 | 9.2 |

**Statistical Methods**

**A) Influence of genetic drift.** Prior to drawing inferences on the magnitude of selection from observed changes in allele frequencies, it is essential to first evaluate whether random genetic drift can be ruled out as a potential influential source. To evaluate the magnitude of genetic drift required to account for the observed temporal variation in allele frequencies, we used the methods-of-moments estimator developed by Nei and Tajima (1981), Pollak (1983), and Waples (1989), which assumes the use of effectively neutral markers. The key indicator variable for estimating $N_e$ is

$$\phi_{ijk} = \widehat{F}_{ijk} - \frac{1}{2n_{ij}} - \frac{1}{2n_{ik}}, \tag{A1}$$

where $i$ denotes the nucleotide site, $j$ and $k$ denote two sampling times, and $n_{ij}$ and $n_{ik}$ are the associated numbers of sampled diploid individuals. $\widehat{F}_{ijk}$ is a normalized measure of the observed allele-frequency change between the two time points,

$$\widehat{F}_{ijk} = \frac{(\widehat{p}_{ij} - \widehat{p}_{ik})^2}{\overline{p}_i(1 - \overline{p}_i)}, \tag{A2}$$

where $\widehat{p}_{it}$ is the estimated allele frequency at time $t$, and $\overline{p}_i = (\widehat{p}_{ij} + \widehat{p}_{ik})/2$. The method-of-moments estimator of $N_e$ from two temporal samples is

$$\widehat{N}_e = \frac{T}{2\overline{\phi}}, \tag{A3}$$

where $\overline{\phi}$ is the average value of $\phi_{ijk}$ over all sites in the sample, and $T$ is the number of years seaparating samples (= 1 for samples in adjacent years). (This averaging of the denominator term to obtain a single estimate of $N_e$ rather than obtaining an estimate for each site and then averaging minimizes the spurious sampling variation that can occur with individual ratios). We obtained estimates of the $N_e$ necessary to account for the standardized variance of allele-frequency change across years separately for each suitably polymorphic site on each chromosome in each of the eight pairs of adjacent years, confining such analysis to sites with an average allele frequency $> 0.02$ in the set of samples (and with no 0.0 values).

**Supplemental Table 2.** Estimates of the effective population size necessary to account for observed allele-frequency changes, averaged over all polymorphic sites with minor-allele frequencies $> 0.02$ on a a chromosome-wide basis. Overall means and standard errors are based on the chromosome-specific measures. Results are given for sites with cutoffs for minimum numbers of individuals in a pair of samples of 10, 20, and 40, for time intervals of one and eight years. Each analysis was based on $> 2,000,000$ polymorphic sites, and the 1-year results are averaged over 8 annual intervals. Note that because the $N_e$ estimator deployed is unbiased, negative estimates can arise when the true parametric value is small.

| | 1-year Interval | | | 8-year Interval | | |
|---|---|---|---|---|---|---|
| Chromosome | 10 | 20 | 40 | 10 | 20 | 40 |
| 1 | 230 | 566 | 260 | 332 | 366 | 428 |
| 2 | 130 | 173 | -706 | 618 | 771 | 1210 |
| 3 | 242 | 1243 | -1238 | 37 | 482 | 794 |
| 4 | 96 | 254 | 82 | 2509 | -17355 | -2465 |
| 5 | 1641 | -13 | 264 | 313 | 382 | 603 |
| 6 | 133 | -240 | 187 | 679 | 1095 | 3986 |
| 7 | 135 | 166 | -421 | 763 | 1126 | 3899 |
| 8 | -277 | 1307 | 655 | 1540 | 15161 | -2660 |
| 9 | 107 | 145 | 2170 | 583 | 876 | 2279 |
| 10 | 146 | 217 | -289 | 289 | 361 | 523 |
| 11 | 53 | -48 | -343 | 684 | 1039 | 1669 |
| 12 | 161 | 400 | 44 | 345 | 441 | 620 |
| Mean | 233 | 348 | 55 | 753 | 395 | 907 |
| SE | 133 | 139 | 241 | 188 | 2013 | 588 |

**Estimation of site-specific selection parameters**

**B) Time-averaged selection intensity associated with a site, $\overline{s}$.** The fundamental approach to estimating selection coefficients associated with a site involves simple linear regression against time $t$ of logit-transformed variables (Kimura and Crow 1970; Lynch and Ho 2020), denoted as

$$y_t = \ln[\,p_t/(1-p_t)\,], \tag{B1}$$

where $p_t$ is the minor-allele frequency estimated at time $t$. Because sample sizes and allele frequencies differ among dates, and both influence the sampling variance of $y_t$, we estimated $\overline{s}$ by using weighted least-squares regression. As there are 9 sampling points, time units are denoted below as $t = 0, \ldots, 8$. Note that $y_t$ is undefined if $p_t = 0$ or 1, and here we only employed sampling points for which the the minor-allele frequency (MAF) $> 0.02$ and the sample size $\geq 10$ individuals.

The sampling variance of a regression coefficient, which in this case is an estimate of $\overline{s}$, is minimized by weighting each data point by its information content, which here is equal to the inverse of the sampling variance of $y_t$,

$$w_t = 1/[\,n_t p_t(1-p_t)\,], \tag{B2}$$

where $n_t$ is the twice the number of diploid individuals in the sample at time $t$ (Lynch 1987). This shows that the optimal weights differ according to allele frequency even if sample sizes are constant.

From Lynch and Walsh (1998), Equation 8.36b, the weighted estimate of the average selection coefficient at the site is

$$\overline{s} = \frac{\sum_{t=0}^{8} w_t(y_t - \overline{y}_w)(t - \overline{t}_w)}{\sum_{t=0}^{8} w_t(t - \overline{t}_w)^2} = \frac{\overline{yt_w} - (\overline{y}_w \cdot \overline{t}_w)}{\overline{t_w^2} - \overline{t}_w^2}, \tag{B3a}$$

where the components are defined as

$$\overline{y}_w = \sum_{t=0}^{8} w_t y_t/w, \quad \overline{y_w^2} = \sum_{t=0}^{8} w_t y_t^2/w, \quad \overline{t}_w = \sum_{t=0}^{8} w_t t/w, \quad \overline{t_w^2} = \sum_{t=0}^{8} w_t t^2/w,$$

$$\overline{yt_w} = \sum_{t=0}^{8} w_t y_t t/w, \quad w = \sum_{t=0}^{8} w_t, \tag{B3b}$$

where the summation is over a number smaller than 9 if there are missing data. Equation B3a is simply equal to the ratio of the weighted covariance and the variance of the weighted times.

From Lynch and Walsh (1998), Equation 8.37b, the sampling variance of $\overline{s}$ is estimated as

$$\sigma^2(\overline{s}) = \frac{\sigma_e^2}{(T-2)(\overline{t_w^2} - \overline{t}_w^2)}, \tag{B4}$$

where

$$\sigma_e^2 = \sum_{t=0}^{8} w_t(\,y_t - a - \overline{s}t\,)^2/w, \tag{B5a}$$

is the weighted residual variance,

$$a = \overline{y}_w - \overline{s}\overline{t}_w \tag{B5b}$$

is the intercept of the regression, and $T$ is the number of samples in the analysis ($T = 9$ if data are present for each year). The square root of Equation B4 is the standard error of the estimate $\overline{s}$.

**Temporal variance of $s$ within sites.** To evaluate the degree to which selection intensity may be fluctuating through time, we estimated the variance of $s$ among single-year time intervals, $\sigma_s^2$, at each site. Because estimates of $s$ from adjacent intervals share the sample at the intermediate time point, these must be avoided in the estimation of $\sigma_s^2$. With a set of 8 consecutive estimates of $s$ at a site, this can be accomplished by performing two sets of analyses, the first involving $s_{0,1}$, $s_{2,3}$, $s_{4,5}$, and $s_{6,7}$, and the second involving $s_{1,2}$, $s_{3,4}$, $s_{5,6}$, and $s_{7,8}$.

The fundamental measures in the following analyses are the estimated interval-specific selection coefficients based on the ratios of logit transforms,

$$s_{i,i+1} = \ln\left(\frac{(1-p_i)p_{i+1}}{p_i(1-p_{i+1})}\right), \tag{B6}$$

where $p_i$ and $p_{i+1}$ are the estimated allele frequencies at time points $i$ and $i+1$. From Equation 12 in Lynch (1987), the sampling variance of this measure is

$$\mathrm{Var}(s_{i,i+1}) \simeq \left(\frac{1}{n_i p_i(1-p_i)} + \frac{1}{n_{i+1}p_{i+1}(1-p_{i+1})}\right), \tag{B7}$$

where $n_i$ is twice the number of individuals sampled at time $i$.

The estimated variance of true selection coefficients is equal to the difference between the variance among the estimates of $s$ and their average sampling variance. For each of the two sets of interval-specific $s$ estimates, the weighted estimate of the variance of $s$ among time intervals is obtained as

$$\widehat{\sigma}_s^2 = \left[ \frac{T}{T-1} \left( \overline{s^2} - \tilde{s}^2 \right) \right] - \mathrm{Var}(s), \tag{B8}$$

where $T$ is the number of samples used in the estimate (here we confine the analyses to sets with $T = 3$ or 4). For the first set of samples,

$$\overline{s^2} = \sum_{i=0}^{3} w_{2i} s_{2i,2i+1}^2 / w, \quad \tilde{s} = \sum_{i=0}^{3} w_{2i} s_{2i,2i+1} / w, \quad \mathrm{Var}(s) = \sum_{i=0}^{3} w_{2i} \mathrm{Var}(s_{2i,2i+1}) / w, \tag{B9a}$$

where the weights are defined as the reciprocal of Equation B7, and $w$ is the sum of the weights used in the summation. (Note that because $w_{2i} = 1/\mathrm{Var}(s_{2i,2i+1})$, $\mathrm{Var}(s)$ is equivalent to the harmonic mean of the individual sampling variances). For the second set of samples,

$$\overline{s^2} = \sum_{i=0}^{3} w_{2i+1} s_{2i+1,2i+2}^2 / w, \; \tilde{s} = \sum_{i=0}^{3} w_{2i+1} s_{2i+1,2i+2} / w, \; \mathrm{Var}(s) = \sum_{i=0}^{3} w_{2i+1} \mathrm{Var}(s_{2i+1,2i+2}) / w. \tag{B9b}$$

The final methods-of-moment estimator of $\sigma_s^2$ for a site is taken to be the average of the two separate estimates.

**Temporal covariance of site-specific selection coefficients.** The focus here is on evaluating whether selection coefficients at individual nucleotide sites are correlated across different time intervals, using Equations B6 and B9 as the units of analysis. The time interval between pairs of estimated selection coefficients to be used in the covariance analysis is denoted as $T$, with $T = 1$ denoting pairs of selection coefficients separated by an entire time interval, e.g., $s_{0,1}$ paired with $s_{2,3}$, $s_{1,2}$ with $s_{3,4}$, etc. Comparison of intervals involving shared intermediate sampling points (i.e., $T = 0$) is not pursued in order to avoid the very substantial nonindependence issues involving adjacent $s$ based on $p_{i-1}$, $p_i$, and $p_{i+1}$, i.e., if $p_i$ is overestimated $s_{i-1,i}$ will be upwardly biased whereas $s_{i,i+1}$ will be correspondingly downwardly biased, causing a bias towards negative covariances. In this particular study, there are nine sampling times, labeled as $i = 0, \ldots, 8$, enabling estimation of the within-site covariance of $s$ for $T = 1$, 2, 3, and 4, and hence to evaluate the time scale over which selection covariance (if any) decays. The specific details for these four $T$ are covered below sequentially.

For any set of $N$ pairs of interval-specific $s$, the sample covariance has the usual definition,

$$\mathrm{Cov}(x, y) = \frac{N(\overline{xy} - \overline{x} \cdot \overline{y})}{N-1}, \tag{B10a}$$

where an overline denotes an average value. The sampling variance of $\mathrm{Cov}(x, y)$ is

$$\mathrm{Var}[\,\mathrm{Cov}(x,y)\,] = \frac{\mathrm{Var}(x)\mathrm{Var}(y) + \mathrm{Cov}^2(x,y)}{N}, \tag{B10b}$$

from Lynch and Walsh (1998; Equation A1.14), and the standard error of $\mathrm{Cov}(x, y)$ is estimated by the square root of Equation B10b. Likewise, the variance of variable $x$ is defined as

$$\mathrm{Var}(x) = \frac{N(\overline{x^2} - \overline{x}^2)}{N-1}, \tag{B11a}$$

which has an estimated sampling variance of

$$\mathrm{Var}[\,\mathrm{Var}(x)\,] = \frac{2\mathrm{Var}^2(x)}{N}. \tag{B11b}$$

Parallel expressions follow for $\mathrm{Var}(y)$ and $\mathrm{Var}[\,\mathrm{Var}(y)\,]$.

$\mathbf{T} = 1$. Depending on data availability, the covariance can be based on up to six pairs of selection coefficients: $s_{0,1}$ and $s_{2,3}$; $s_{1,2}$ and $s_{3,4}$; $s_{2,3}$ and $s_{4,5}$; $s_{3,4}$ and $s_{5,6}$; $s_{4,5}$ and $s_{6,7}$; and $s_{5,6}$ and $s_{7,8}$. Here, we required there to be at least four pairs of data to estimate site-specific parameters. An issue with this full $T = 1$ analysis is the presence of slight bias (sampling scheme 1 in Lynch and Ho 2020). Although the members of each $(x, y)$ pair are based on different samples, some of the selection coefficients within the set of $x$ are also contained within the set of $y$. The expected covariance owing to shared samples is

$$-[(N/(N-1)] \cdot E[\,\overline{x} \cdot \overline{y}\,] = -\,1/[N(N-1)] \cdot E[\,s_{1,2}s_{2,3} + s_{2,3}s_{3,4} + s_{3,4}s_{4,5} + s_{4,5}s_{5,6} + s_{5,6}s_{6,7} + s_{2,3}^2 + s_{3,4}^2 + s_{4,5}^2 + s_{5,6}^2\,],$$

where only terms with nonzero expectations (i.e., containing shared samples) are shown, and in this case with the full set of data $N = 6$. The set of five cross-product terms share samples from time-points, $i = 2$, 3, 4, 5, and 6 respectively, and from Lynch and Ho (2020; Equation A8a) have expectations equal to $-1/(n_i p_i(1 - p_i)$, where $n_i$ denotes the number of sampled haplotypes (twice the number of diploid individuals); the terms are negative because the shared frequency is in the numerator of one $s$ and the denominator of the other, Equation B6. The four squared terms, indexed with $i, j$ have excess expectations $[1/(n_i p_i(1 - p_i)] + [1/(n_j p_j(1 - p_j)]$. After collecting terms, the bias is found to be

$$B_c = -\frac{1}{N(N-1)} \cdot \sum_{i=3}^{5} \frac{1}{n_i p_i(1-p_i)} = -\frac{1}{N(N-1)} \cdot \sum_{i=3}^{5} w_i, \tag{B12a}$$

where $w_i$ is as defined in Equation B2. As the bias is negative, the original estimate must be increased by the same amount, i.e., the bias-free estimator is

$$\text{Cov}(x, y)_c = \text{Cov}(x, y) - B_c. \tag{B12b}$$

Note that with the above sampling scheme, if data are missing, $N$ must accordingly be reduced. Equation B12a still applies, except for two cases: if $s_{1,2}$ is missing, $w_2$ must be included in the summation of Equation B12a; and if $s_{6,7}$ is missing, $w_6$ must be included in the summation of Equation B12a.

In contrast, the use of overlapping data points results in upward bias of the estimated temporal variance of $s$. For the first set of estimates (in the set of $x = s_{0,1}$, $s_{1,2}$, $s_{2,3}$, $s_{3,4}$, $s_{4,5}$, and $s_{5,6}$), the bias in the variance is

$$B_{vx} = \frac{2}{N(N-1)} \sum_{i=1}^{5} w_i, \tag{B13a}$$

and the unbiased estimator is obtained by modifying Equation B11a to

$$\text{Var}(x)_c = \text{Var}(x) - B_{vx}. \tag{B13b}$$

If there are missing $s$ values, $N$ must be reduced accordingly, and terms must be removed from Equation B13a according to the subscripts on the missing data. For example, if $s_{0,1}$ is missing, the first term in B13a is eliminated (as subscript 1 is involved). If $s_{1,2}$ is missing, the first two terms in B13a are eliminated (subscripts 1 and 2). If $s_{1,2}$ and $s_{3,4}$ are missing, $N = 4$, all but the final term in B13a is removed (subscripts 1, 2, 3, and 4).

For the second set of estimates (in the set of $y = s_{2,3}$, $s_{3,4}$, $s_{4,5}$, $s_{5,6}$, $s_{6,7}$, and $s_{7,8}$),

$$B_{vy} = \frac{2}{N(N-1)} \sum_{i=3}^{7} w_i, \tag{B14a}$$

yielding the unbiased estimator by modifying Equation B11a to

$$\text{Var}(y)_c = \text{Var}(y) - B_{vy}. \tag{B14b}$$

For cases in which data are missing, the same rules noted above apply: $N$ is reduced accordingly; and terms are removed from $B_{vy}$ where subscripts correspond with the missing data.

Finally, two other $T = 1$ analyses can be performed with reduced sets of data, sharing no sampling dates between $x$ and $y$ sets: the first uses $x = s_{0,1}$, $y = s_{2,3}$ and $x = s_{4,5}$, $y = s_{6,7}$;

and the second uses $x = s_{1,2}$, $y = s_{3,4}$ and $x = s_{5,6}$, $y = s_{7,8}$. An average of the results from the two analyses then provides overall unbiased estimates of the covariances and variances. In this case, the sampling variance of the covariance is simply equal to the square root of the variance of the two estimates, substituting these for $x$ in Equation B11a, and dividing by $\sqrt{2}$.

**T** = 2. Here, the possible entry pairs become: $s_{0,1}$ and $s_{3,4}$; $s_{1,2}$ and $s_{4,5}$; $s_{2,3}$ and $s_{5,6}$; $s_{3,4}$ and $s_{6,7}$; and $s_{4,5}$ and $s_{7,8}$. The bias for the covariance with a full set of observations ($N = 5$) is:

$$B_c = -\frac{w_4}{N(N-1)}. \tag{B15}$$

If $s_{2,3}$ is missing, $w_3$ must be added to the numerator; if $s_{3,4}$ is missing, $(w_3 + w_4)$ must be added to the numerator; if $s_{4,5}$ is missing, $(w_4 + w_5)$ must be added to the numerator; and if $s_{5,6}$ is missing, $w_5$ must be added to the numerator.

The bias for the variance of $x$ for a full set of observations is

$$B_{vx} = \frac{2}{N(N-1)} \sum_{i=1}^{4} w_i, \tag{B16}$$

$N$ is reduced accordingly if data pairs are missing. If $s_{0,1}$ is missing, $w_1$ is removed from the summation; if $s_{1,2}$ is missing, $(w_1 + w_2)$ is removed; if $s_{2,3}$ is missing, $(w_2 + w_3)$ is removed; if $s_{3,4}$ is missing, $(w_3 + w_4)$ is removed; if $s_{4,5}$ is missing, $w_4$ is removed.

The bias for the variance of $y$ for a full set of observations is

$$B_{vy} = \frac{2}{N(N-1)} \sum_{i=4}^{7} w_i, \tag{B17}$$

Again, $N$ is reduced accordingly if data pairs are missing. If $s_{3,4}$ is missing, $w_4$ is removed from the summation; if $s_{4,5}$ is missing, $(w_4 + w_5)$ is removed; if $s_{5,6}$ is missing, $(w_5 + w_6)$ is removed; if $s_{6,7}$ is missing, $(w_6 + w_7)$ is removed; if $s_{7,8}$ is missing, $w_7$ is removed.

All of the other expressions noted above for $T = 1$ hold, after substituting the above expressions for the biases to obtain corrected estimates of the variances and covariances.

Again, two other $T = 2$ analyses can be performed with reduced sets of data, sharing no sampling dates between $x$ and $y$ sets, and eliminating any need for bias correction: the first uses $x = s_{0,1}$, $y = s_{3,4}$ and $x = s_{1,2}$, $y = s_{4,5}$; and the second uses $x = s_{3,4}$, $y = s_{6,7}$ and $x = s_{4,5}$, $y = s_{7,8}$. (In neither set is there a shared date between $x$ and $y$ variables). An average of the two then provides an overall unbiased estimate.

**T** $= 3$. Here, the possible entry pairs become: $s_{0,1}$ and $s_{4,5}$; $s_{1,2}$ and $s_{5,6}$; $s_{2,3}$ and $s_{6,7}$; and $s_{3,4}$ and $s_{7,8}$, and we required all four pairs to be available. For a full set of data ($N = 4$), the bias in the covariance is:

$$B_c = +\frac{w_4}{N(N-1)}. \tag{B18}$$

Note that the bias is positive in this case.

The bias for the variance of $x$ for a full set of observations is

$$B_{vx} = \frac{2}{N(N-1)} \sum_{i=1}^{3} w_i, \tag{B19}$$

The bias for the variance of $y$ for a full set of observations is

$$B_{vy} = \frac{2}{N(N-1)} \sum_{i=5}^{7} w_i, \tag{B20}$$

Here, there is a single unbiased, reduced set of data, sharing no sampling dates between $x$ and $y$ sets: $x = s_{0,1}$, $y = s_{4,5}$ and $x = s_{2,3}$, $y = s_{6,7}$.

**T** $= 4$. Here, the possible data pairs become: $s_{0,1}$ and $s_{5,6}$; $s_{1,2}$ and $s_{6,7}$; and $s_{2,3}$ and $s_{7,8}$. In this case, there is no bias in the covariance estimator, as the $x$ and $y$ sets do not share sampling dates.

**C) Genome-wide covariance of selection across time intervals.** All of the approaches outlined above focus on the temporal dynamics of variants at individual nucleotides sites in order to evaluate site-specific magnitudes of selection and the degree to which these vary and covary across time. An alternative approach to extracting information from a temporal series of allele frequencies is to consider the aggregate behavior of all allele-frequency changes across specific time intervals (Buffalo and Coop 2019, 2020; Lynch and Ho 2020). The key unit of measurement here is

$$\Delta_{i,t} = \widehat{p}_{i,t} - \widehat{p}_{i,t-1}, \tag{C1}$$

the change in MAF estimates for nucleotide site $i$ across the time interval $t-1, t$.

The goal is to determine whether there is nonzero covariance between allele-frequency changes for the full set of $N$ genomic sites, using

$$\mathrm{Cov}(\Delta_t, \Delta_{t'}) = \frac{\sum_{i=1}^{N} (w_{i,t}\Delta_{i,t}) \cdot (w_{i,t'}\Delta_{i,t'})}{\sum_{i=1}^{N} w_{i,t} \cdot w_{i,t'}} - \overline{\Delta}_t \cdot \overline{\Delta}_{t'}, \tag{C2a}$$

where to minimize the sampling variance, we use the weights

$$w_{i,t} = \left( \frac{\widehat{p}_{i,t}(1-\widehat{p}_{i,t})}{n_{i,t}} + \frac{\widehat{p}_{i,t-1}(1-\widehat{p}_{i,t-1})}{n_{i,t-1}} \right)^{-1}, \tag{C2b}$$

where the sample sizes in the denominators are equal to twice the number of individuals, and

$$\overline{\Delta}_t = \frac{\sum_{i=1}^N w_{i,t}\Delta_{i,t}}{\sum_{i=1}^N w_{i,t}}. \tag{C2c}$$

Note that if $t' = t+1$ (adjacent time intervals), Equation C2a is downwardly biased by the shared intermediate time point, which induces negative sampling covariance. In this case, an unbiased estimate can be obtained by adding the sampling variance of allele frequency for the intermediate time point. Using Equation C2b to obtain the inverse of the weights, the weighted sampling variance is estimated as

$$\text{Var}(\widehat{p}_t) = \frac{\sum_{i=1}^N n_i^2/[\widehat{p}_{i,t}(1-\widehat{p}_{i,t})]}{\sum_{i=1}^N n_i^3/[\widehat{p}_{i,t}(1-\widehat{p}_{i,t})]^2}, \tag{C2d}$$

which reduces to the expected $p(1-p)/n$ with constant allele frequencies and sample sizes.

The standard errors of $\text{Cov}(\Delta_t, \Delta_{t'})$ are obtained by use of the square root of Equation B10b, with the covariance term defined by Equation C2a, and the variance terms defined by the univariate analogs,

$$\text{Var}(\Delta_t) = \frac{\sum_{i=1}^N w_{i,t}\Delta_{i,t}^2}{\sum_{i=1}^N w_{i,t}} - \overline{\Delta}_t^2. \tag{C2e}$$

### D) Computer Simulations of Fluctuating Selection

To determine the effects of fluctuating selection on the long-term average within-population diversity, we employed the computational machinery described in Lynch (2020). This is a straight-forward "Wright-Fisher" population-genetic framework with discrete generations, fixed absolute population sizes, and biallelic loci. Each generation involves consecutive episodes of mutation, selection, and random genetic drift. Mutation is bidirectional, and selection operates in an additive manner. Here, the simulations involve a single pair of loci, one neutral and the other subject to Gaussian fluctuations in the selection intensity on one of the alleles at the selected site each generation, with mean $\overline{s}$, variance $\sigma_s^2$, and no temporal correlation. The simulations are run long enough, after an ample burn-in period, to obtain stable estimates of the mean heterozygosity at each locus. The code is written so that multiple populations with different sizes can be simultaneously evaluated in parallel.

**E) Potential Effects of an Egg Bank on Parameter Estimates**

The study population is an intermittent pond that dries up annually (generally by June) and remains so until the following spring. Given the extreme drying conditions, we do not expect much survival of resting eggs beyond a single year, it cannot be entirely ruled out either. We, therefore, pursued computer simulations of allele-frequency dynamics under the assumption of some degree of viable resting-egg retention from year to year.

To consider the consequences of a potential egg bank on short-term allele-frequency fluctuations and the resultant estimates of $N_e$, a simple Wright-Fisher model of random genetic drift was produced, with each annual sample drawing from the pool of hatching resting eggs, followed by random drift based on an annual effective population size of $N_e$. A fraction $h$ of resting eggs was assumed to hatch each year, leading to a geometric decline in contribution over the subsequent $x$ years of $h(1-h)^x$. This is an extreme view of an egg-bank effect, as it assumes indefinite viability until eventual hatching. For each value of $h$ (ranging from 1.0 to 0.5) and a full spectrum of starting allele frequencies (ranging from 0.05 to 0.50), $> 10^7$ estimates of $N_e$ were obtained from temporal strings of allele-frequency data using the methods outlined above, after allowing the population to equilibrate for a long enough time period for all eggs in the earliest cohort to hatch, and assuming no sampling error (i.e., $n = \infty$). The program used for these simulations, NeBias.cpp, is available at https://github.com/LynchLab/Daphnia-egg-bank-models-Drift-Effects. As shown in Supplemental Figure 1, egg-bank retention leads to downward bias in estimates of annual $N_e$, but the effect is quite small, $< 20\%$ downward bias even with egg-retention rates as high as 50%. The ratio of observed $N_e$ relative to its actual annual value is very close to 1.0 for $h > 0.95$, and for lower $h$ is closely approximated by

$$\frac{N_{obs}}{N_e} \simeq h \left( 1 + 1.57 \sum_{i=1}^{\infty} (1-h)^i \right). \tag{E1}$$

To consider the consequences of an egg bank for estimates of the mean and temporal standard deviation of selection intensity ($\overline{s}$ and $\sigma_s$) , similar simulations were performed, but assuming zero stochasticity associated with random genetic drift and sampling so as to investigate bias effects associated with temporal selection alone. As in the actual field study, nine annual samples were used to derive each pair of parameter estimates, and the entire procedure was replicated $\sim 10^9$ times to compare average estimates with their rue expected values. The program used for these simulations, SelBias.cpp, is available at https://github.com/LynchLab/Daphnia-egg-bank-models-Selection-Estimation-Effects.

As shown in Supplemental Figure 2, egg-bank retention results in slight upward bias in estimates of $\sigma_s$, but never by more than 15%. The effects on estimates of $\overline{s}$ are more pronounced. Provided $\sigma_s \leq 0.01$, the absolute value of $\overline{s}$ is downwardly biased by up to two-fold (for the extreme situation of 70% annual resting-egg survival); in this case, provided $\sigma_s/\overline{s} \leq 10$, the fractional depression in estimates of $\overline{s}$ is closely approximated by $(1-h)$, i.e., 20% resting-egg retention results in a 20|% downward bias. For $10 \geq \sigma_s/\overline{s} \leq 100$, the downward bias is slightly greater, but unlikely to exceed 70%; there is, however, intrinsic downward bias of the estimator of $\overline{s}$ in this case $\simeq$ 50%, even in the absence of an egg bank. For $\sigma_s/\overline{s} \geq 1000$, which requires very small $\overline{s}$ and/or very large $\sigma_s$, the bias can be substantially larger, even shifting the sign of the average $\overline{s}$ estimate. Note, however, that the absolute deviations in $\overline{s}$ estimates remains small in these cases.

**Supplemental Figure 1.** Reduction in estimates of effective population sizes based on allele-frequency estimates from pairs of adjacent annual samples made at the time of resting-egg hatching, relative to the actual effective size at the annual time of sexual reproduction, with a fraction $h$ of resting eggs assumed to hatch each year, with a fraction $(1 - h)$ assumed to survive each subsequent year until all have eventually hatched. The results are essentially independent of the actual effective size and of the minor-allele frequency (not shown, with averages across all frequencies plotted in the figure). The curved line is given by Equation A4.

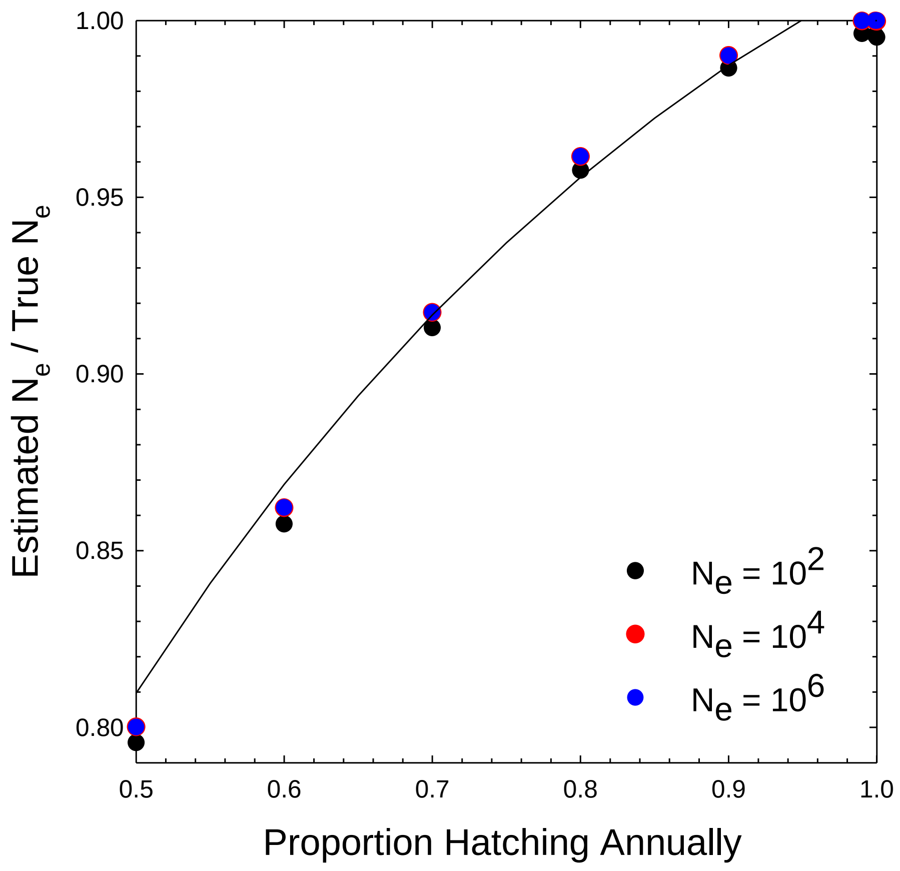

**Supplemental Figure 2.** Effects of the presence of an egg bank on average estimates of selection coefficients and their temporal standard deviation ($\overline{s}$ and $\sigma_s$), based on simulated data under the assumption that $s$ is Gaussian distributed, with negligible effects of random genetic drift (as supported by the analyses herein, for short-term changes) and no sampling error on the part of the investigator (to evaluate the true evolutionary consequences). A fraction $h$ of resting eggs are assumed to hatch each year, with a fraction $(1 - h)$ surviving each subsequent year until all have eventually hatched. Closed symbols denote results for $\overline{s}$, whereas open symbols denote $\sigma_s$ estimates. Black triangles denote results for $\overline{s}$ when $h = 1.0$.

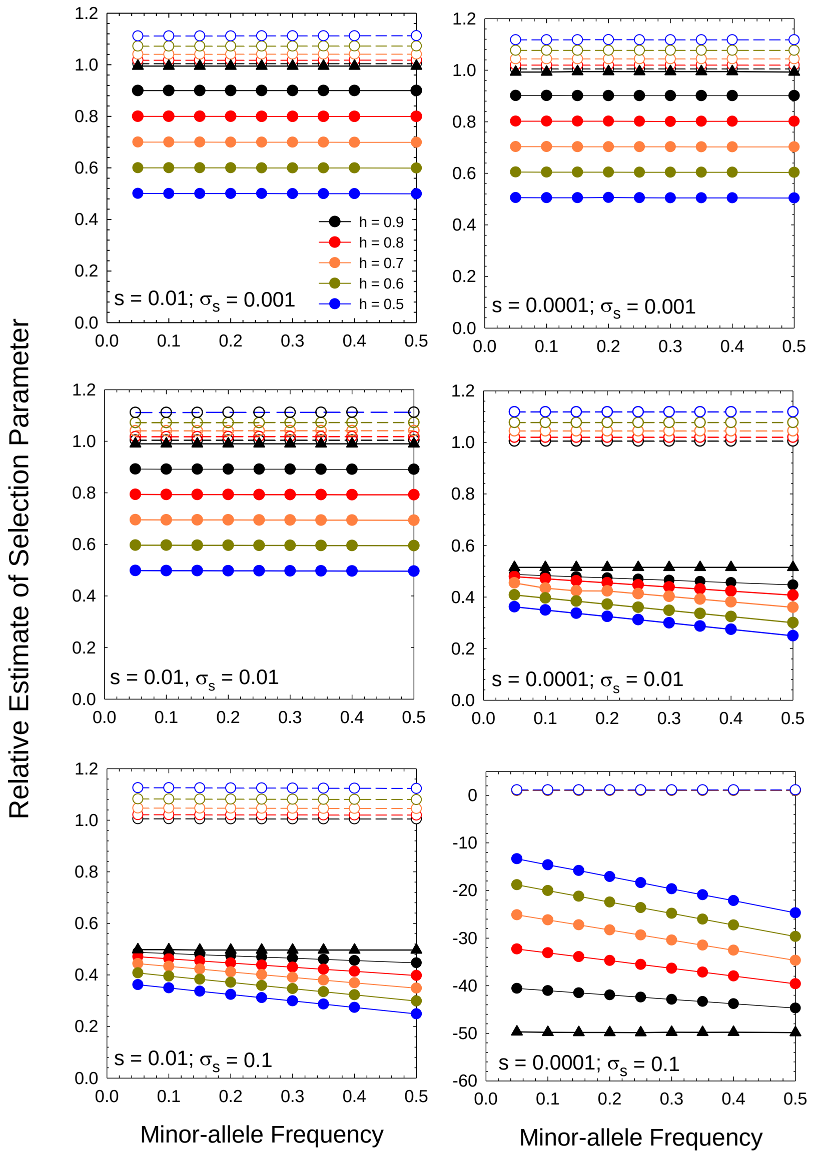
